# Isolation and Screening of Polyethylene Terephthalate (PET)-degrading bacteria from Municipal Waste Disposal Site in West Java, Indonesia

**DOI:** 10.1101/2025.06.16.659912

**Authors:** Theresia Evelyn, Anissa Fauziah Zulfitri, Azzania Fibriani

## Abstract

Polyethylene terephthalate (PET) is a persistent plastic contributing to environmental pollution due to its resistance to natural degradation. This study aimed to identify PET-degrading bacteria from two major landfills in West Java, that is Bantar Gebang and Cipayung. Samples were collected from soil and plastic waste at three locations per site. A combined approach was used: metagenomic analysis to profile microbial diversity, and culturable enrichment using PET-supplemented mineral medium followed by isolation and PEG-based clear zone screening. PET degradation was assessed over 21 days using PET film as the sole carbon source, and surface damage was evaluated via scanning electron microscopy (SEM). Metagenomic analysis revealed high microbial diversity dominated by *Pseudomonadota* and *Bacillota*, suggesting potential for PET degradation. From 34 culturable isolates, six exhibited clear zone activity: BP8, CT9, CT11 (Bantar Gebang) and ET4, EP19, FP20 (Cipayung). SEM and relative enzyme activity assays confirmed PET film degradation, particularly in isolates BP8 and EP19. Molecular identification via 16S rRNA sequencing revealed high similarity of BP8 to *Priestia megaterium*, CT9 to *Comamonas terrae*, CT11 and FP20 to *Brucella pseudintermedia*, ET4 to *Shinella yambaruensis*, and EP19 to *Micrococcus luteus*. These results highlight the potential of landfill-derived bacteria for PET biodegradation and offer candidates for further biotechnological development in plastic waste management.

**Importance:** Plastic pollution is a growing problem, especially in landfills where it can remain for decades without breaking down. This study looked for bacteria in two major landfills in West Java, Indonesia, that might help degrade PET, a common plastic used in bottles and packaging, using a combination of two scientific approaches: metagenomics (analyzing microbial DNA directly from the environment) and culturable methods (growing microbes in the lab). By combining different scientific methods, we were able to find several types of bacteria that show early signs of being able to break down PET in the lab. While further studies are needed to confirm how well these microbes work and how they break down plastic, these initial findings provide a useful starting point for developing biological methods to manage plastic waste more sustainably in the future.

## 1. Introduction

Plastics have become an integral part of daily life due to their lightweight, durability, affordability, and strength. However, their resistance to natural degradation has led to significant plastic waste accumulation in the environment. Among synthetic polymers, polyethylene terephthalate (PET) stands out due to its superior mechanical, chemical, and thermal properties, attributable to its chemical structure consisting of polar ester groups and aromatic rings derived from terephthalic acid (TPA). Commercial PET products such as beverage bottles and biaxially oriented films typically exhibit high crystallinity, around 30–40%, high melting points (255–265°C), and varying glass transition temperatures (67°C for amorphous PET, ∼80°C for semi-crystalline PET) [1],[2]. However, these favourable industrial characteristics simultaneously contribute to PET’s resistance to natural biodegradation, resulting in prolonged environmental persistence. Indeed, PET waste significantly exacerbates global plastic pollution, with accumulation projected to persist for centuries [3].

Indonesia is one of the largest contributors to plastic waste globally. According to the National Solid Waste Management Information System [4], Indonesia generated approximately 58.1 million tons of waste annually in 2024, with plastics constituting approximately 19.56% of this total. Unfortunately, most low-value plastics, including PET, predominantly end up unmanaged in landfills rather than being recycled [4]. Conventional methods like incineration and mechanical recycling face significant limitations, such as high energy demands, pollutant emissions, and economic inefficiencies, underscoring the need for sustainable alternatives [5], [6].

The enzymatic degradation of PET by microbes is increasingly recognized as an effective strategy for addressing plastic pollution in an environmentally sustainable way. A variety of microorganisms are known to break down PET through the synthesis of specific enzymes. For example, *Ideonella sakaiensis*, a bacterium discovered at a plastic recycling site, is known to secrete PETase and MHETase, enzymes that work together to depolymerize PET into its basic monomers [7]. Likewise, genera such as Thermobifida [8], Bacillus [9], Pseudomonas [10], Brucella (formerly Ochrobactrum) [11], Stenotrophomonas, Comamonas [12], and Saccharomonospora [13] have also shown significant PET degradation potential.

However, research on the isolation and utilization of local bacteria from plastic-contaminated environments in Indonesia remains limited. Given the large volume of plastic waste in Indonesia, this study aims to isolate and identify PET-degrading bacteria from environments heavily exposed to plastic pollution and evaluate their potential for PET biodegradation in vitro. Bantar Gebang and Cipayung landfills, two of the largest and most plastic-polluted sites in Indonesia, were selected as sampling locations. Bantar Gebang landfill is located in Bekasi, West Java, covering a total area of 110.3 hectares. In 2023, the total area of these landfills had expanded to 117.5 hectares. According to the Jakarta Environmental Agency, this landfill is the largest in Asia. Bantar Gebang landfill has been in operation since 1989, serving as the primary facility for managing waste from all residents of DKI Jakarta. Each day, this site receives approximately 6,500 to 7,000 tons of waste from DKI Jakarta [4]. Cipayung landfill is located in Depok, West Java, with a total area of 11,6 hectares. Cipayung landfill has been in operation since 1994. This landfill accommodates waste from 1/3 of the total area of Depok City, receiving a daily waste volume of approximately 1300 tons [14]. The large amount of waste received has caused the landfill to reach its full capacity. Among the total waste composition at Cipayung landfill, plastic waste for the largest portion, comprising 16.66% within the category of recyclable inorganic waste [15]. These two locations serve as highly strategic research sites for exploring bacteria capable of degrading plastic, particularly PET, which can be utilized in biological PET waste management efforts. This is supported by the high volume of plastic waste and the environmental conditions of both landfill sites.

This study explains how one kind of plastic, PET (polyethylene terephthalate) can be degraded by bacteria or also called biodegradation. Biodegradation of plastic is defined as a decomposition process that utilizes the activity of microorganisms so that changes occur in plastic [16]. Plastic biodegradation generally proceeds through multiple stages. It begins with biodeterioration, where environmental factors and microbial activity work together to break down plastic into smaller fragments, and microorganisms start forming biofilms on the surface. This is followed by depolymerization, in which extracellular enzymes secreted by microbes cleave the long polymer chains into shorter oligomers and oxidize them, forming chemical groups such as carbonyl and hydroxyl. Next is the assimilation stage, where these degradation products are absorbed into microbial cells through membrane receptors and undergo internal metabolism. Finally, during mineralization, these compounds are fully oxidized and converted into simpler end-products, including carbon dioxide (CO_2_), methane (CH_4_), water (H_2_O), and mineral salts, thus completing the biodegradation process [17].

## 2. Materials and Methods

### 2.1 Sample Collection

Soil and plastic waste samples were collected randomly from the three points selected of the dumpsite. This resulted in 3 soil samples and 3 plastic waste samples collected from each landfill, Bantar Gebang landfill and Cipayung landfill. The samples were collected randomly from the superficial layer of soil (5-20 cm) in depth, using pre-sterilized spatula and were transferred into sterile centrifuge tube. Samples stored at 4°C for microbiological analysis and -20°C for metagenomic. In the laboratory, soil samples from different collection points were pooled before proceeding with DNA extraction and sequencing.

### 2.2 DNA Extraction and Metagenomic Sequencing

DNA was extracted using the ZymoBIOMICS DNA Kit following the manufacturer’s instructions. Quality and quantity were assessed via agarose gel electrophoresis and a Nanodrop spectrophotometer. DNA libraries were prepared using the TruSeq DNA Library Preparation Kit v2 (Illumina) and quantified with a Qubit fluorometer. Sequencing was conducted at the Bandung Institute of Technology using an Illumina HiSeq 2500 platform.

### 2.3 Media Preparation for PET Degrading Bacteria

For bacterial enrichment and screening, Mineral Salts Medium (MSM) was used with various supplements:

- MSM broth with 0.1% PET solution for enrichment and degradation testing.
- MSM broth with PEG and glucose for initial degradation assessments.
- MSM agar with 0.2% PEG for clear zone screening.
- PET agar for isolation and purification.

For enrichment culture of PET degrading bacteria, a Mineral Salts medium (MSM) containing polyethylene terephthalate (PET) was prepared by 5 g K_2_HPO_4_, 2 g NaH_2_PO_4_, 4 g (NH_4_)_2_SO_4_, 0,5 g MgSO_4_.7H_2_O, 0,5 g NaCl, 0,15 g KCl, and 0,2 g CaCl_2_.6H_2_O. In addition, the medium contained 1 mL mixed trace element solution (0,1 g each FeSO_4_.7H_2_O, ZnCl_2_, and CuSO_4_ in 100 mL distilled water), and 1 mL Tween 80 as a surfactant agent. These ingredients were dissolved in 1 L distilled water (pH was fixed at 7,2) in schott duran bottle and then media was sterilized in an autoclave at 121 °C for 15 min [18]. The Mineral Salts medium was supplemented with 0,1% (w/v) PET solution. PET solution was prepared by small pieces of PET bottle that previously had been crushed and disinfected 30 min in 70% ethanol and air-dried for 30 min in laminar air flow chamber with exposure to UV light. A small pieces of crushed PET sheet were added to 10 ml DMSO at about 0,1 g/ml in a heat-stable borosilicate glass (100 ml volume). The glass was covered with aluminium foil and heated to about 180 °C. PET solution was stirred with a magnetic stirrer until the PET was dissolved [19]. The PET solution was added to the sterile Mineral Salts medium. For determining clear zone formation around the colony, a Mineral Salts medium supplemented with 0,2% (w/v) PEG and 1,5% (w/v) agar [20].

PET agar for isolation and purification PET degrading bacteria consist two layer: the bottom layer and top layer. The bottom layer consist of 50% R2A medium, was prepared by 9,1 g of R2A powder and 6,0 g bacteriological agar in 1 L distilled water. The top layer containing PET consist of 15,5 g/L bacteriological agar, the anionic surfactant Sarkosyl at 0,1 g/L were added to 1x PBS (pH 7,4) and supplemented with 0,1 % (w/v) PET solution [19].

### 2.4 Enrichment Culture, Isolation, and Purification of PET Degrading Bacteria

One gram of each soil sample was suspended in 50 mL sterile physiological saline (0.85% NaCl), shaken at 120 rpm for 4 hours, and 5 mL of the supernatant was inoculated into 100 mL MSM containing PET. After one week of incubation at 30°C, subculturing was repeated into fresh PET-supplemented MSM for two enrichment cycles. Cultures were then spread on PET agar and nutrient agar, incubated at 30°C for 3–10 days. Single colonies were isolated, purified, and activated on PET agar for downstream assays.

### 2.5 Screening of the Best Isolates for PET Biodegradation Assay

Isolates were screened on PEG-supplemented MSM agar by clear zone formation. After two weeks at 30°C, plates were stained with 0.1% Coomassie Brilliant Blue R-250 and decolorized using 40% methanol with 10% acetic acid to visualize enzymatic degradation halos [21]. Clear zone ratios were measured using ImageJ software and then calculated as clear zone ratio = (Total area of isolate + clear zone − Area of isolate) / Area of isolate.

### 2.6 Sanger Sequencing and Phylogenetics Analysis

The inoculum was initially propagated in liquid Luria Bertani (LB) medium and incubated at 37°C for 24 hours with shaking at 150 rpm. Subsequently, total genomic DNA was extracted from pure single colonies cultured on Nutrient Agar (NA) using the ZymoBIOMICS DNA Miniprep Kit (Zymo Research, USA), following the manufacturer’s protocol. The purity and concentration of the extracted DNA were assessed using a Nanodrop spectrophotometer (Thermo Scientific, USA). Verified DNA samples served as templates for amplifying the 16S rRNA gene using universal primers: forward primer 27F (5′-AGAGTTTGATCCTGGCTCAG-3′) and reverse primer 1492R (5′-GGTTACCTTGTTACGACTT-3′) [18]. PCR products were subsequently examined via agarose gel electrophoresis to confirm successful amplification and sent to an external facility for Sanger sequencing. Sequencing data were quality-trimmed, contigs were assembled, and phylogenetic trees were constructed using Geneious Prime software.

### 2.7 Microscopic Analysis

Microscopic analysis was performed to determine the Gram reaction and morphology of the isolates. A single colony from PET-agar was heat-fixed on a slide and subjected to Gram staining using crystal violet, iodine, ethanol (decolorizer), and safranin. The stained slides were examined under a light microscope (1000×, oil immersion). Additionally, cell morphology (shape, size, and arrangement) was observed and recorded for each isolate.

### 2.8 Biochemical Tests

Biochemical tests were performed on 24-hour-old bacterial cultures grown on PET agar at 37°C to characterize the metabolic and enzymatic properties of the isolates. The catalase test was conducted by adding hydrogen peroxide to the bacterial culture to observe bubble formation, indicating the presence of the catalase enzyme. The ability to ferment glucose, lactose, and sucrose was assessed using Triple Sugar Iron Agar (TSIA) medium. Additionally, indole production, motility, and hydrogen sulfide (H_2_S) production were evaluated using SIM (Sulfide-Indole-Motility) medium. The Methyl Red/Voges-Proskauer (MR-VP) test was performed on facultative anaerobic isolates to determine their sugar fermentation pathways. The results from these biochemical tests provided insights into the metabolic capabilities of the PET-degrading bacterial isolates.

### 2.9 Biodegradation Studies

A biodegradation test was performed using of PET films (600 mm x 600 mm x 0,35 mm, Goodfellow) were cut by hole puncher and the final size of PET film is about 6 mm-diameter. PET films had been disinfected 30 min in 70% ethanol and air-dried for 30 min in laminar air flow chamber with exposure to UV light. The film were aseptically added to erlenmeyer flask containing 100 ml of sterile mineral salts medium. Each flask was inoculated with 5 ml of 24 h old culture grown in mineral salts medium supplemented with 0,1% glucose. After inoculate, cultures were incubated on a orbital shaker at room temperature and 120 rpm for 7-21 days. Flask without inoculation served as sterile control. Every 7 days interval, bacterial growth and pH changed was measured.

### 2.10 Topography Analysis Using Scanning Electron Microscopic (SEM)

Surface topography analysis of PET film was conducted before and after the biodegradation assay. Post-biodegradation analysis was performed on PET films exposed to the three bacterial isolates that exhibited the highest biodegradation activity, as determined by the clear zone assay. Two approaches were used for topographical analysis: bacterial attachment observation without SDS treatment and surface examination after SDS treatment.

For surface examination, PET films were washed sequentially with 2% sodium dodecyl sulfate (SDS), ethanol, and distilled water for 30 minutes each, followed by air drying [22]. The dried films were then fixed, vacuum-dried, and coated with a gold/palladium layer before being scanned using Scanning Electron Microscopy (SEM) coupled with Energy Dispersive X-ray Spectroscopy (EDS) (JEOL NeoScope JCM-7000) at 15 kV.

### 2.11 Enzyme Activity Assay

Extracellular proteins were precipitated using pre-chilled sterile acetone at a 4:1 volume ratio. The mixture was vortexed, incubated at −20 °C for 3 h, and centrifuged at 13,000–15,000 × g for 10 min. The supernatant was discarded, and residual acetone was evaporated at room temperature. The protein pellet was resuspended in 1 mL PBS and stored at −20 °C. PETase activity was measured using p-nitrophenyl butyrate (pNPB) as the substrate in 96-well plates. Each well contained 200 µL resuspended protein and 2 µL pNPB. Absorbance was recorded at 405 nm using a GloMax® Discover System. While no calibration curve was applied, absorbance values were used to indicate relative enzymatic activity between isolates.

### 2.12 Statistical Analysis

Biodegradation assay data, including clear zone diameter and associated activity, were statistically analyzed using SPSS software.

## 3. Results and Discussions

### 3.1 Unculturable Study: Metagenomic Analysis

The Bantar Gebang and Cipayung landfills (**Figure 1**) are two of the largest waste disposal sites in Jakarta, Indonesia. These sites have been operational for more than 20 years and have accumulated significant amounts of municipal solid waste over time. The accumulation of plastic waste, in particular, remains a significant concern, as it poses long-term environmental and ecological risks. In this study, metagenomic analysis was employed to profile the microbial communities associated with soil and plastic waste samples from both locations. This approach enabled the detection of both culturable and non-culturable microbes, emphasized the value of metagenomics in identifying microbial and enzymatic diversity involved in plastic degradation [23]. Beside that, plastisphere microbial communities differ significantly from bulk soil, often harboring specialized degraders [24]. Also, this non-culturable approach enables comprehensive profiling of microbial taxa, including non-cultivable or slow-growing organisms that are often overlooked in traditional isolation methods.

**Figure 1.**
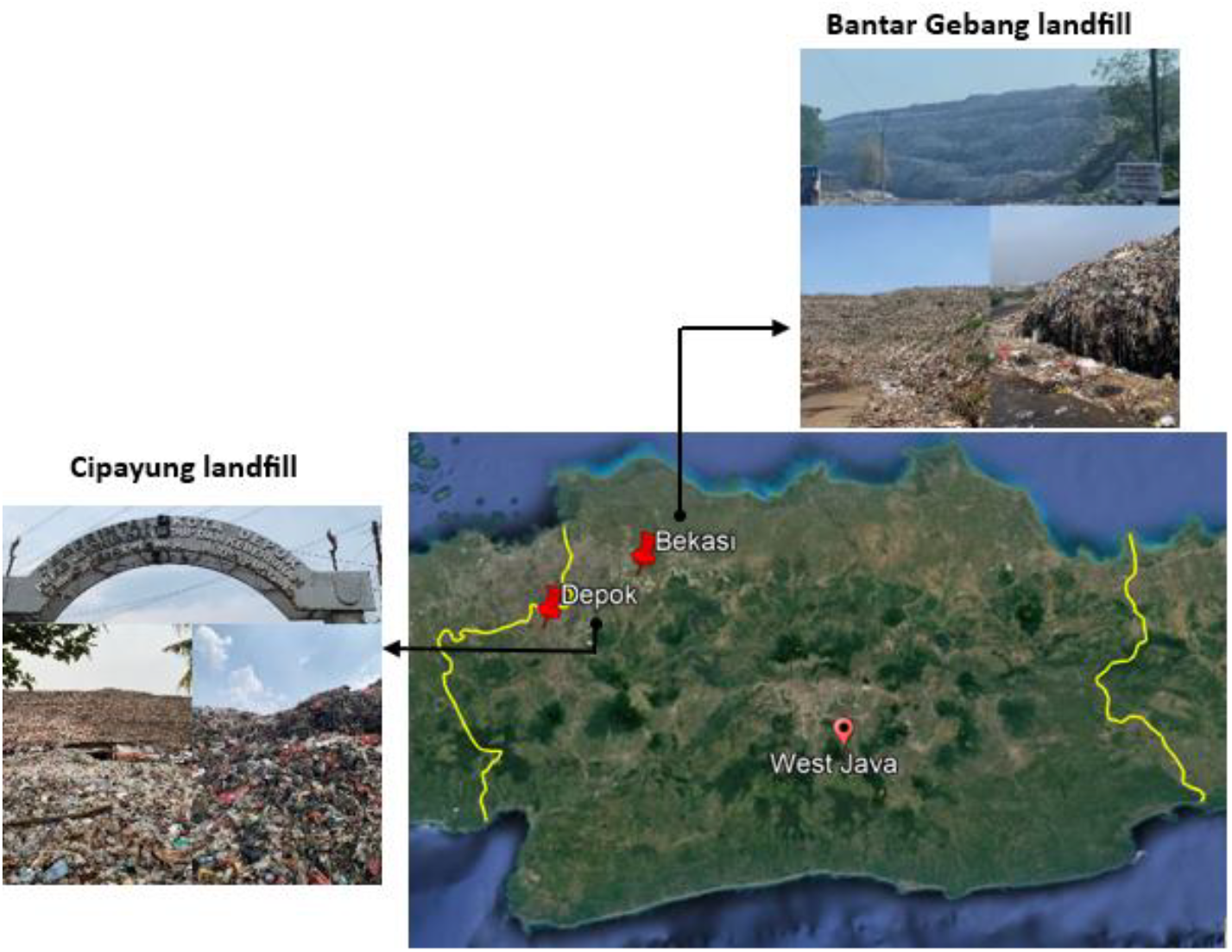
Sampling site: Bantar Gebang and Cipayung Landfill in Bekasi and Depok, West Java.

The metagenomic analysis provided insight into the microbial community composition in soil and plastic-associated samples from the two landfill sites, Bantar Gebang (CP and CT) and Cipayung (EP and ET), based on sequencing depth and data robustness. Microbial diversity, evaluated through the Shannon diversity index, indicated that all samples harbored diverse communities. Notably, EP from Cipayung exhibited the highest diversity (Shannon index = 7.25), whereas CP from Bantar Gebang also showed significant diversity (Shannon index = 6.02). Richness estimates paralleled these findings, with CT having the highest OTU richness (6491), closely followed by CP (6301). Phylum-level relative abundance (>0.5% cutoff) highlighted distinct microbial composition patterns (**Figure 2**).

**Figure 2.**
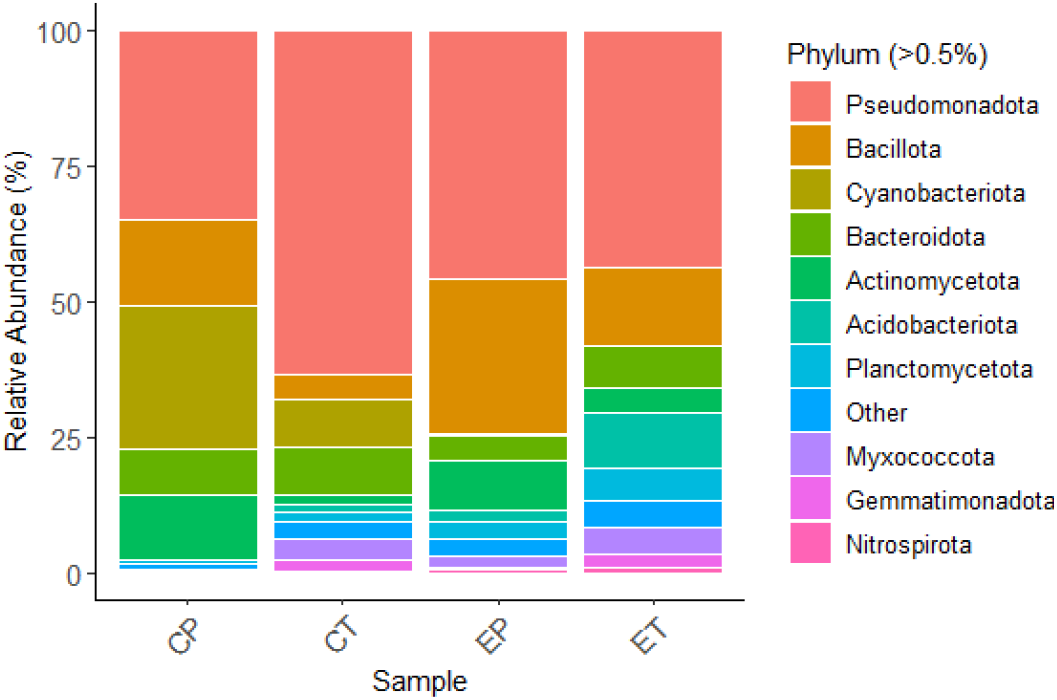
Microbial community composition at the phylum level based on relative abundance (>0.5% cutoff, others grouped as “Other”) from soil and plastic waste samples collected from two landfills. CP and CT were collected from Bantar Gebang landfill, EP and ET were collected from Cipayung landfill.

A key ecological trend emerged: Bacillota was enriched in plastic-associated samples (CP and EP), indicating a possible selective advantage in colonizing plastic substrates. This aligns with Bacillota’s metabolic versatility and tolerance to xenobiotics. Conversely, soil-derived sa, ples (CT and ET) exhibited more taxonomic balance across phyla including Actinomycetota and Acidobacteriota, consistent with the higher nutrient and structural complexity of bulk soil environments. Interestingly, even though ET had lower sequencing depth (**Table 2**), its Shannon index (6.17) remained comparable, underscoring the role of substrate type in structuring microbial communities. These community shifts suggest that prolonged plastic exposure may act as an ecological filter, enriching taxa capable of hydrocarbon utilization or biofilm-based colonization, potentially including PET-degrading candidates.

**Table 1.** Sample collection points and characteristics.

| Sampling site | Sample | Coordinate | Sample Appearance |
| --- | --- | --- | --- |
| Bantar Gebang landfill | A | 6°21'26.15"S<br>106°59'45.69"E | Brown soil, dry and compact, mixed waste (plastic, fabric, other materials) |
|  | B | 6°21'25.78"S<br>106°59'49.29"E | Black soil, soft and wet, dominated by plastic waste (surface and buried) |
|  | C | 6°21'18.31"S<br>107°00'02.31"E | Brown soil, dry and compact, dominated by buried fabric and plastic waste reaching the surface |
| Cipayung landfill | E | 6°25'16.19"S<br>106°47'19.28"E | Brown soil, wet material, dominated by plastic waste |
|  | F | 6°25'15.35"S<br>106°47'19.18"E | Dry soil with a dark appearance, dominated by plastic and fabric waste |
|  | G | 6°25'16.87"S<br>106°47'19.51"E | Dry soil with a dark appearance and plant debris |

**Table 2.** Diversity index of microbial community.

| Sam-<br>ple | Reads | Pielou's<br>evenness | Rich<br>ness | Shannon<br>diversity |
| --- | --- | --- | --- | --- |
| CP | 303246 | 0.69 | 6301 | 6.02 |
| CT | 225154 | 0.77 | 6491 | 6.76 |
| EP | 122504 | 0.83 | 6198 | 7.25 |
| ET | 3554 | 0.79 | 1038 | 6.17 |

### 3.2 Culturable Study: Isolation, Purification, and Screening of PET Degrading Bacteria

A total of 12 environmental samples underwent a three-stage enrichment process using minimal media with PET powder as the sole carbon source. From subsequent isolation on PET-agar through serial subculturing and purification, 21 isolates were retained and subjected to functional screening based on their ability to form clear zones on PEG-infused agar plates (**Figure 3**). The clear zone assay is based on the hydrolytic activity of enzymes secreted by bacteria that degrade polyethylene glycol (PEG), a polymer analog used here to simulate PET ester linkages. Out of 21 isolates, 14 demonstrated detectable clear zones, and 3 isolates showed higher degradation from each sapling points (**Table 3**). The three most promising isolates from Bantar Gebang were BP8 (0.344 ± 0.102), CT9 (0.220 ± 0.063), and CT11 (0.165 ± 0.048), while EP19 (0.779 ± 0.104), FP20 (0.579 ± 0.139), and ET4 (0.400 ± 0.067) were the top performers from Cipayung.

**Table 3.** Clear zone ratio formed by each isolate.

| Isolate | Isolate growth area<br>(cm<sup>2</sup>) | Clear zone area<br>(cm<sup>2</sup>) | Ratio ± Stdev |
| --- | --- | --- | --- |
| **Bantargebang Landfill** |  |  |  |
| BP8 | 0,815 | 0,469 | 0,578±0,033<sup>a</sup> |
| CT11 | 3,610 | 0,553 | 0,166±0,047<sup>bc</sup> |
| CT9 | 0,545 | 0,117 | 0,220±0,063<sup>b</sup> |
| BP7 | 1,494 | 0,165 | 0,113±0,024<sup>bc</sup> |
| BT6 | 1,099 | 0,104 | 0,085±0,026<sup>c</sup> |
| CP12 | 0,619 | 0,059 | 0,095±0,021<sup>c</sup> |
| BT5 | 0,649 | 0,042 | 0,065±0,019<sup>c</sup> |
| **Cipayung Landfill** |  |  |  |
| EP19 | 0,670 | 0,522 | 0,779±0,104<sup>a</sup> |
| FP20 | 0,720 | 0,313 | 0,435±0,139<sup>b</sup> |
| ET4 | 0,802 | 0,307 | 0,400±0,067<sup>bc</sup> |
| FT8 | 0,359 | 0,108 | 0,307±0,075<sup>cd</sup> |
| FT14 | 0,685 | 0,129 | 0,191±0,047<sup>d</sup> |
| ET2 | 0,265 | 0,056 | 0,209±0,015<sup>d</sup> |
| EP17 | 0,558 | 0,084 | 0,151±0,036<sup>d</sup> |
| FT13 | 0,528 | 0,115 | 0,204±0,084<sup>d</sup> |
| FT14 | 0,685 | 0,129 | 0,191±0,047<sup>d</sup> |

Mean differences are indicated by lowercase superscript letters. Statistical comparisons were conducted separately for each sampling location. Values are presented as mean ± standard deviation (SD) from four replicates.

**Figure 3.**
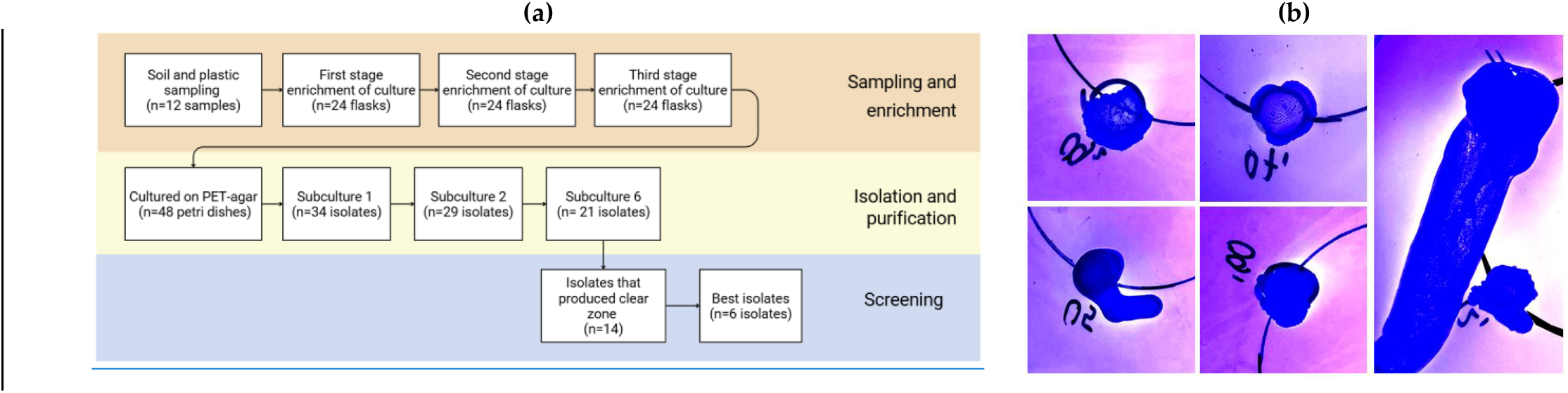
(a) Process of selecting the best isolates for the biodegradation test with PET film. (b) Representative images of clear zones formed by selected bacterial isolates grown on minimal medium supplemented with polyethylene glycol (PEG) as the sole carbon source. Plates were incubated for 7 days at 37 °C and stained with Coomassie Brilliant Blue. Clear zone around colonies indicates enzymatic hydrolysis of PEG, as a proxy for polyesterase activity.

### 3.3 Isolate Characterization

To complement molecular identification, macroscopic, microscopic, and biochemical characterizations were performed for the six isolates selected based on their clear zone assay. 16S rRNA gene sequencing was performed to identify the isolates (**Table 5**). The isolate BP8, showed 99.9% similarity to *Priestia megaterium* ATCC 14581, a known member of the Firmicutes phylum with documented metabolic versatility. CT11 was closely related (99.4%) to *Comamonas terrae*, a member of the Proteobacteria phylum known for its role in environmental pollutant degradation. CT9 and FP20 were both affiliated with *Brucella pseudintermedia*, sharing 98.8% and 98.7% sequence identity, respectively. Interestingly, isolate EP19 matched *Micrococcus luteus* (99.8%), an Actinobacteria, while ET4 aligned closely with *Shinella yambaruensis* (99.7%), an Alphaproteobacteria species. These taxonomic assignments demonstrate the phylogenetic diversity of PET-degrading bacteria isolated from landfill environments.

To contextualize the taxonomic affiliations of the isolated strains, a phylogenetic tree was constructed using 16S rRNA sequences, aligned via MUSCLE and inferred using the maximum likelihood (PhyML) method with 100 bootstrap replicates (**Figure 4**). *Bacteroides fragilis* was selected as the outgroup, anchoring the phylogeny and validating the evolutionary trajectory of the isolates. The tree delineates the phylogenetic diversity of the isolates, which span across three major bacterial phyla: Bacillota (e.g., *Priestia megaterium*), Actinomycetota (e.g., *Micrococcus luteus*), and Pseudomonadota (e.g., *Comamonas terrae, Shinella yambaruensis*, and *Brucella pseudintermedia*). The robustness of the tree topology was supported by high bootstrap values across key nodes, affirming the taxonomic resolution provided by 16S rRNA gene sequencing.

**Figure 4.**
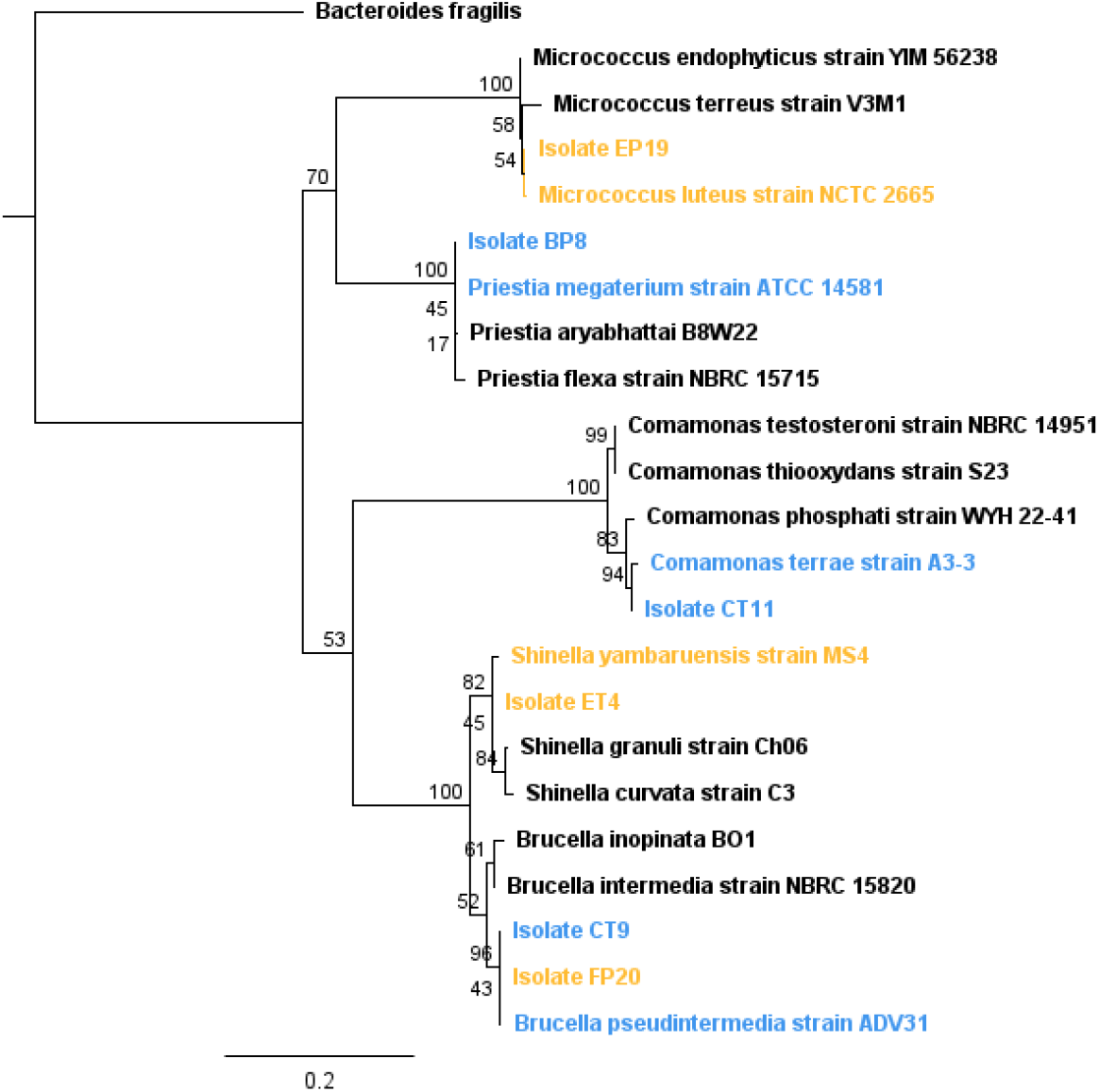
Maximum likelihood phylogenetic tree showing the taxonomic relationships of six PET-degrading bacterial isolates (highlighted in blue) based on 16S rRNA gene sequences. The tree was constructed using the PhyML algorithm with 100 bootstrap replicates and multiple sequence alignment performed via MUSCLE in Geneious Prime. Bacteroides fragilis was used as the outgroup, selected from the same family but a different genus to root the tree.

Interestingly, isolates CT9 and FP20 clustered closely with *Brucella pseudintermedia*, sharing over 98.7% sequence identity and forming a well-supported clade, indicative of close evolutionary lineage and possibly similar functional roles. Likewise, CT11 aligned with *Comamonas terrae*, a genus well-known for its ability to degrade environmental pollutants, including recalcitrant plastics. Isolate EP19 showed high similarity to *Micrococcus luteus*, a member of Actinomycetota with reported capabilities in surface colonization and oxidative metabolism—traits relevant for PET degradation. The presence of *Priestia megaterium* (BP8), a Gram-positive bacillus with broad enzymatic versatility, and *Shinella yambaruensis* (ET4), an Alphaproteobacteria typically found in soil.

Colonies exhibited distinct morphological features consistent with their taxonomic identities (**Table 4**). BP8 and ET4 formed irregular, white colonies typical of Gram-negative bacilli, whereas CT11 displayed smooth, glossy colonies and were Gram-positive coccobacilli. EP19 was distinct in producing yellow-pigmented colonies and was identified as Gram-positive cocci, aligning with its identity as *Micrococcus luteus*. CT9 and EP20, associated with *Brucella pseudintermedia*, formed small, white colonies and stained as Gram-negative cocci.

**Table 4.**
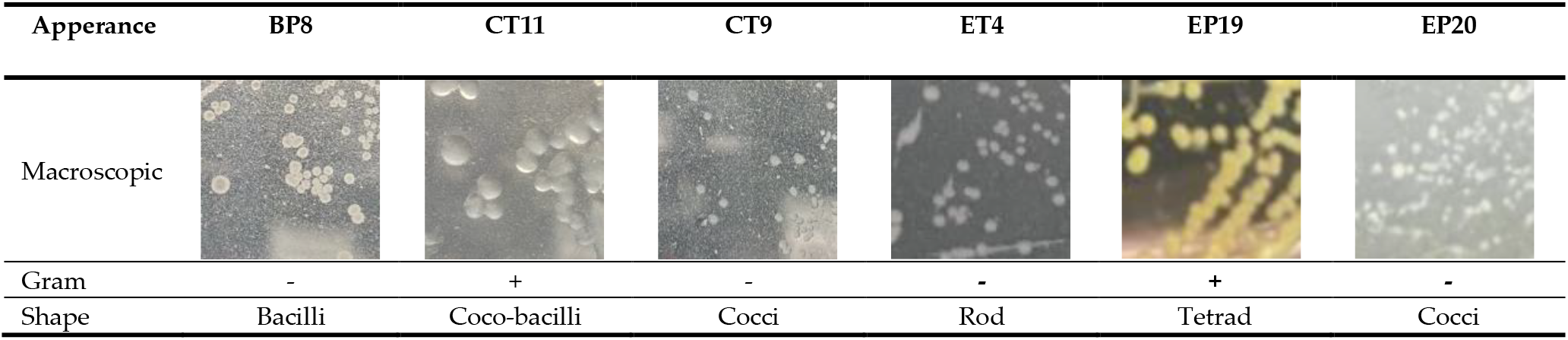
Macroscopic and microscopic morpholgy of isolates.

| Apperance | BP8 | CT11 | CT9 | ET4 | EP19 | EP20 |
| --- | --- | --- | --- | --- | --- | --- |
| Macroscopic |  |  |  |  |  |  |
| Gram | - | + | - | - | + | - |
| Shape | Bacilli | Coco-bacilli | Cocci | Rod | Tetrad | Cocci |

**Table 5.** Isolate identification based on 16s rRNA as marker gene with NCBI database.

| Isolate code | % Similarity | Closest species |
| --- | --- | --- |
| BP8 | 99,9 | *Priestia megaterium* ATCC 14581 |
| CT11 | 99,4 | *Comamonas terrae* A3-3 |
| CT9 | 98,8 | *Brucella pseudintermedia* ADV31 |
| ET4 | 99,7 | *Shinella yambaruensis* MS4 |
| EP19 | 99,8 | *Micrococcus luteus* NCTC 2665 |
| FP20 | 98,7 | *Brucella pseudintermedia* ADV31 |

All six isolates demonstrated catalase activity (**Table 6**), a trait commonly observed in aerobic environmental bacteria and particularly relevant to plastic biodegradation. This enzyme serves as a protective mechanism against the oxidative stress associated with polymer breakdown. During the biodegradation of synthetic plastics such as polyethylene terephthalate (PET), oxidative reactions lead to the generation of reactive oxygen species (ROS), including hydrogen peroxide. Catalase plays a crucial role in mitigating this stress by catalyzing the decomposition of hydrogen peroxide into water and oxygen, thereby preventing cellular damage and ensuring bacterial viability in plastic-rich environments [25]. Beyond its classical antioxidant function, recent multi-omics studies have suggested that catalase and catalase-peroxidase enzymes may also participate directly in oxidative depolymerization pathways, facilitating the initial cleavage of plastic polymers [26].

**Table 6.** Biochemical test of isolates.

| Apperance | BP8 | *Priestia megaterium* | CT11 | *Comamonas* sp. | CT9 | *Brucella* sp. | ET4 | *Shinella yambaruensis* | EP19 | *Micrococcus luteus* | FP20 | *Brucella pseudintermedia* |
| --- | --- | --- | --- | --- | --- | --- | --- | --- | --- | --- | --- | --- |
| Sulfide production | - | - | - | ? | + | - | - | - | - | - | - | - |
| Indole production | - | - | - | - | - | - | - | - | - | - | - | - |
| Ferment glucose | + | + | - | - | - | - | - | - | - | - | - | - |
| Ferment sucrose | - | - | - | - | - | - | - | - | - | - | - | - |
| Ferment lactose | - | - | - | - | - | - | - | - | - | - | - | - |
| Catalase | + | - | + | + | + | + | + | + | + | + | + | + |
| Motility | + | + | + | + | + | + | - | - | - | - | + | + |

- : negative ? : unclear or not determined
+ : positive n: no source

Furthermore, the non-fermentative profile of all isolates and their consistent inability to metabolize common sugars anaerobically reinforces their classification as obligate aerobes, in agreement with their expected environmental niches. This metabolic profile is advantageous in PET degradation, as aerobic respiration is typically coupled with extracellular enzyme secretion and oxidative attack on polymeric substrates. Motility was also detected in most isolates, potentially enhancing their ability to colonize plastic surfaces, a key initial step in biofilm formation and subsequent biodegradation.

### 3.4 Biodegradation Test with PET Film

The assessment of enzyme activity (**Figure 5**) and PET film surface damage by scanning electron microscopy (SEM) (**Table 7**) highlights the biodegradation potential of six selected bacterial isolates from two major landfills in Indonesia. Among these, isolate BP8 from Bantargebang demonstrated the highest enzymatic activity within its group and substantial PET surface disruption under SEM. *Priestia megaterium* is well known for its ability to form robust biofilms and produce extracellular enzymes, including esterases and lipases, which are crucial for the hydrolytic breakdown of polyester-based plastics [27]. The presence of pits and cracks on PET film treated with BP8 supports its active role in initiating surface depolymerization processes. CT11, affiliated with *Comamonas terrae*, displayed moderate enzyme activity and visible surface erosion, consistent with its taxonomic proximity to *Comamonas testosteroni*, which has been shown to degrade PET via hydrolytic cleavage and assimilate resulting monomers through a series of transporters and intracellular enzymes [28]. In that study, *C. testosteroni* KF-1 demonstrated significant PET fragmentation, generating nanoparticle-sized plastic fragments and exhibiting upregulation of a specific PET hydrolase whose deletion markedly reduced degradation efficiency. The same strain also benefited from acetate supplementation, suggesting a co-metabolic enhancement of PET degradation. Although CT11 may not produce extensive extracellular degradation, its potential for oligomer transport and intracellular bioconversion, facilitated by porin and TPA transporter systems [29], suggests an alternative but equally relevant biodegradation strategy.

**Table 7.**
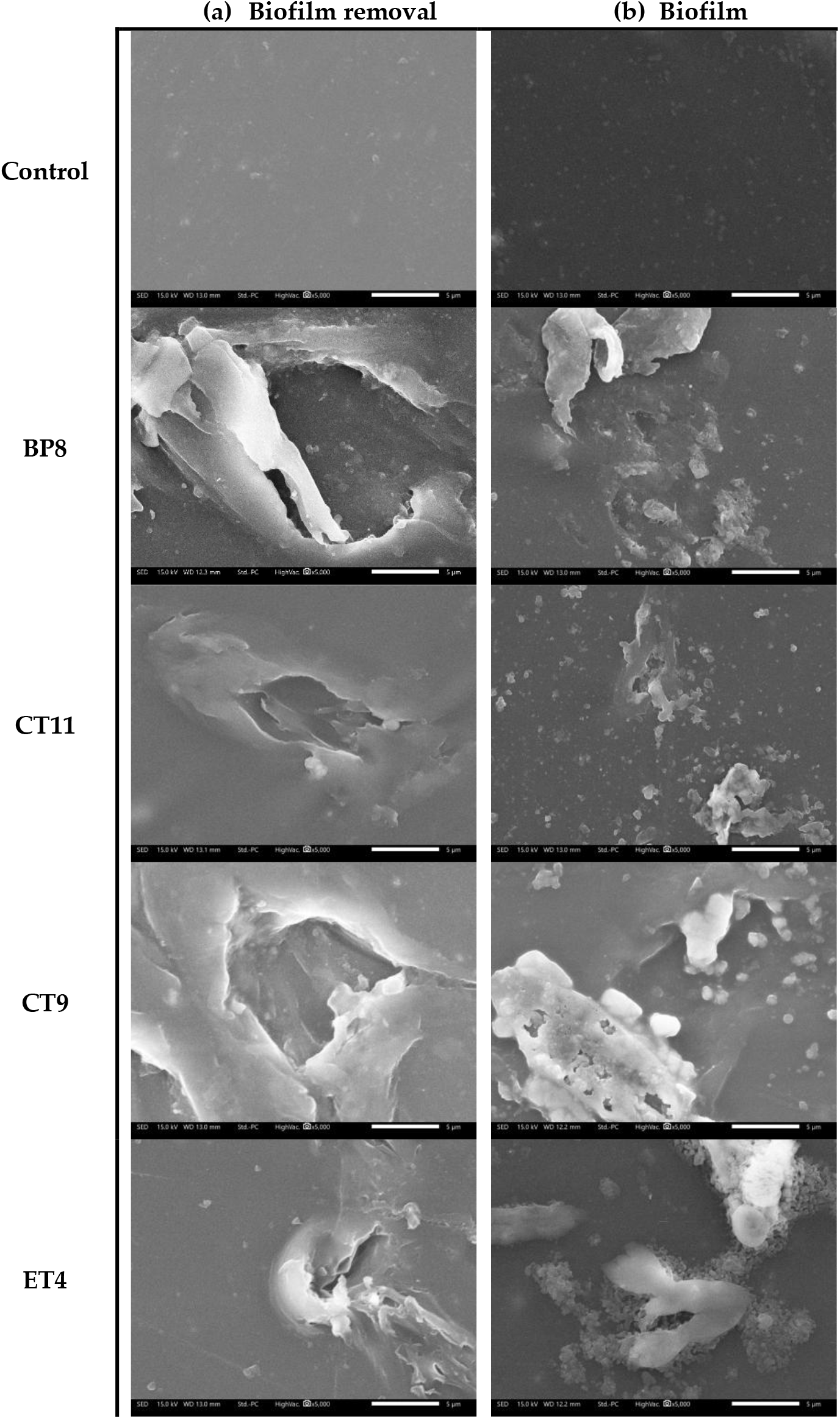
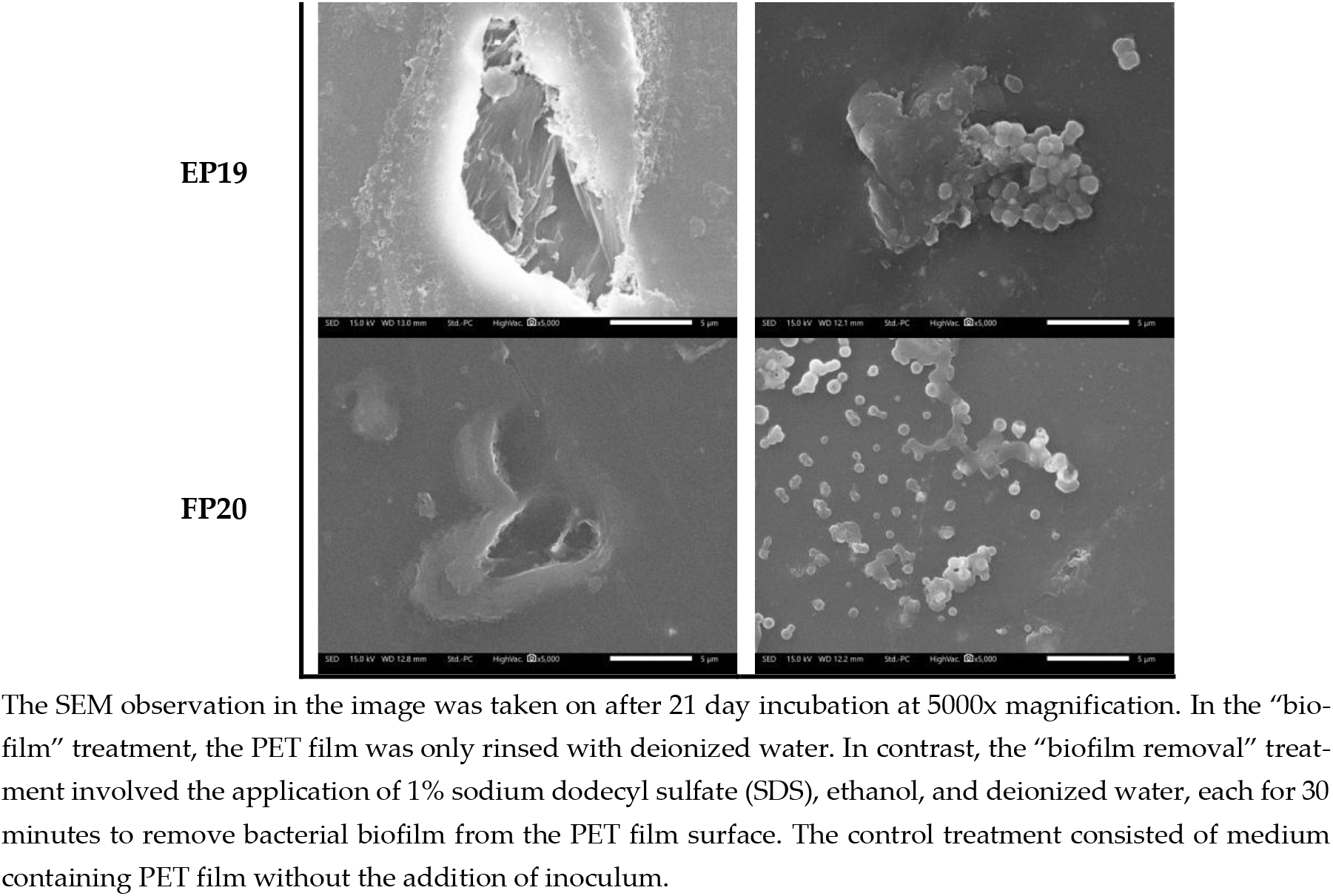
Scanning electron micrograph of PET films after 21-day incubation. (a) SEM analysis revealed PET film surfaces without biofilm, (b) SEM analysis revealed PET film surfaces with biofilm formation.

**Figure 5.**
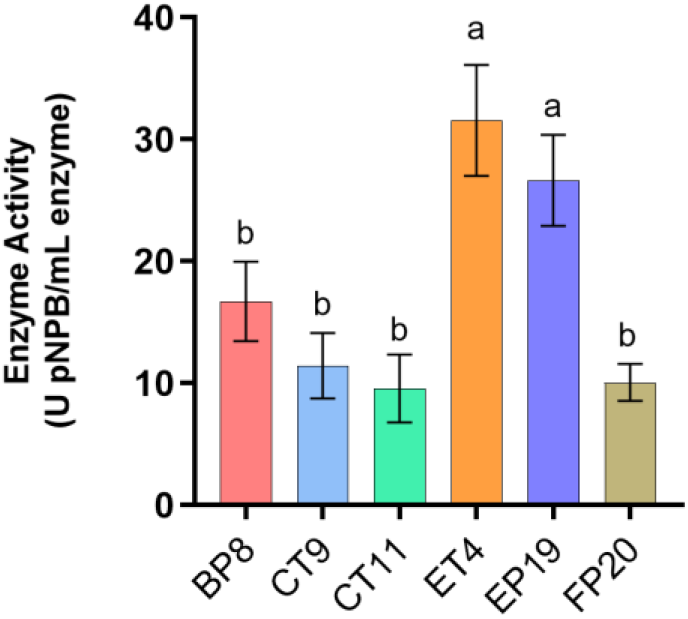
Enzyme activity assay. Crude extracellular proteins from treated isolates were precipitated using acetone and harvested on day 21. The substrate used was p-nitrophenyl butyrate (p-NPB). Absorbance was measured at 405 nm.

From the Cipayung isolates, EP19 exhibited the highest pNPB enzymatic activity and significant PET surface deterioration. This is consistent with previous reports where *Micrococcus luteus* strain CGK112, isolated from cow dung, demonstrated efficient biofilm formation and significant degradation of HDPE, including structural deformation and surface oxidation as observed through SEM, FTIR, and EDX analyses [30]. CT9 and FP20, both identified as *Brucella pseudintermedia*, exhibited relatively low enzymatic activity and modest PET surface disruption under SEM observation. These findings align with previous studies indicating that Brucella-mediated plastic degradation is typically slower and may require extended incubation periods to achieve substantial breakdown. For example, *Brucella intermedia* strain IITR130 achieved only 26.06% PET degradation over a 60-day period [31]. Similarly, recent work demonstrated the involvement of *Brucella pseudintermedia* in polypropylene (PP) microplastic degradation, where degradation rates remained moderate (28.5 ± 0.5%) even after 28 days of incubation [32]. These collective results suggest that while Brucella pseudintermedia may exhibit biodegradative potential, its activity appears to be limited in the early stages and may be more dependent on biofilm formation and hydrolase-mediated processes over prolonged exposure. ET4 (*Shinella yambaruensis*), while exhibiting high enzyme activity, showed less aggressive PET surface disruption. However, its ability to metabolize sulfur-containing compounds may offer an auxiliary advantage in sulfur-rich minimal media environments [33].

## 4. Conclusions

This study successfully identified six PET-degrading bacterial isolates from Bantar Gebang and Cipayung landfills using a combined metagenomic, biochemical profiling, enzymatic assays, and SEM-based approach. Metagenomic analysis revealed diverse microbial communities, with plastic-associated samples (CP and EP) were enriched in Bacillota, suggesting niche adaptation. Functional screening showed PET-degrading potential in all six isolates (BP8, CT9, CT11, ET4, EP19, and FP20) with BP8 and EP19 exhibiting the highest enzymatic activity. SEM analysis confirmed PET surface damage across all isolates, with BP8 and EP19 showing the most significant degradation. The isolates were taxonomically identified via 16S rRNA sequencing as *Priestia megaterium* (BP8), *Comamonas terrae* (CT11), *Brucella pseudintermedia* (CT9 and FP20), *Micrococcus luteus* (EP19), and *Shinella yambaruensis* (ET4). These findings highlight landfill sites as valuable sources of plastic degrading bacteria and support integrated methods for biotechnological applications.

## Author Contributions

All authors contributed equally to this work. All authors have read and agreed to the published version of the manuscript.

## Acknowledgments

We gratefully acknowledge the support of the School of Life Sciences and Technology, Institut Teknologi Bandung, for providing funding and essential research facilities. We also thank the authorities of Bantargebang and Cipayung for granting permission to conduct sampling in their respective areas.

## Disclaimer Note

During the preparation of this work, the author(s) used ChatGPT to assist with drafting, editing, and refining the content. After using this tool, the author(s) reviewed and edited the content as needed and take(s) full responsibility for the content of the publication.

